# Soft selective sweeps predominate in the yellow fever mosquito *Aedes aegypti*

**DOI:** 10.1101/2025.07.06.663360

**Authors:** Remi N. Ketchum, Daniel R. Matute, Daniel R. Schrider

## Abstract

The *Aedes aegypti* mosquito is a vector for human arboviruses and zoonotic diseases and therefore poses a serious threat to public health. Understanding how *Ae. aegypti* adapts to environmental pressures—such as insecticides—is critical for developing effective mitigation strategies. However, most traditional methods for detecting recent positive selection search for signatures of classic “hard” selective sweeps, and to date no studies have examined soft sweeps in *Ae. aegypti.* This is a significant limitation as this is vital information for understanding the pace of adaptation—populations that can immediately respond to new selective pressures are expected to adapt more often via standing variation or recurrent adaptive mutations (both of which may produce soft sweeps) than via *de novo* mutations (which produces hard sweeps). To this end, we used a machine learning method capable of detecting hard and soft sweeps to investigate positive selection in *Ae. aegypti* population samples from Africa and the Americas. Our results reveal that soft sweeps are significantly more common than hard sweeps, which may imply that this species can respond quickly to environmental stressors. This is a particularly concerning finding for vector control methods that aim to eradicate *Ae. aegypti* using insecticides. We highlight genes under selection that include both well-characterized and putatively novel insecticide resistance genes. These findings underscore the importance of using methods capable of detecting and distinguishing hard and soft sweeps, implicate soft sweeps as a major selective mode in *Ae. aegypti,* and highlight genes that may aid in the control of *Ae. aegypti* populations.

## INTRODUCTION

A central goal of evolutionary biology is to uncover the genomic underpinnings of adaptation by identifying loci under positive selection. Researchers have made significant progress in this challenging task in a wide array of species. Some well-studied examples include: the hypoxia pathway gene *EPAS1* that was implicated in differences in hemoglobin concentrations at high altitude in humans (Huerta-Sánchez et al. 2014) or the *Ectodysplasin (EDA)* locus in threespine stickleback fish that facilitated their transition from marine to freshwater (Barrett et al. 2008). These discoveries, as well as many others, have been made possible by theoretical and methodological advances which have allowed for the detection and characterization of selective sweep signatures.

Traditionally, selective sweeps were thought to occur when a *de novo* beneficial mutation appears and then quickly rises in frequency until it becomes fixed in a population (Maynard Smith and Haigh 1974). This type of selective sweep is referred to as a hard sweep and is characterized by a lack of diversity (except that which is introduced by recombination or mutation during the sweep) around the vicinity of the selected site (Maynard Smith and Haigh 1974). A hard sweep may also result in increased linkage disequilibrium (LD) on either side of the sweep (Kelly 1997; Kim and Nielsen 2004), or a skew in allele frequency patterns (Fay and Wu 2000; Nielsen et al. 2005). In contrast, soft sweeps act on a mutation that is initially neutral (or weakly deleterious) and evolves under drift until a change in the selective environment causes the mutation to become beneficial and sweep to fixation (Orr and Betancourt 2001; Hermisson and Pennings 2005). They may also occur when a beneficial allele arises via mutation or migration during the selective phase of a sweep (Pennings and Hermisson 2006a; Pennings and Hermisson 2006b). In soft sweeps, the beneficial allele exists on multiple haplotypes that share a common ancestor prior to the onset of the sweep and so the resulting skew in patterns of genetic diversity around the selected region may be both qualitatively different and less pronounced than that of a hard sweep (Prezeworski et al. 2005; Schrider et al. 2015). There is evidence that in at least some populations, adaptation proceeds mainly via soft sweeps (Garud et al. 2015; Schrider and Kern 2017; Xue et al. 2021). This in turn implies that these populations are able to adapt rapidly, as they need not wait for an adaptive mutation to arise (Karasov et al. 2010). In this way, selective sweep signatures can reveal loci underpinning adaptation as well as how quickly a population can adapt to new stressors, provided one can identify the type of sweep that occurred.

Sweep detection methods, which generally involve calculating single summary and/or test statistics (e.g., (Fay and Wu 2000; Kim and Nielsen 2004; Nielsen et al. 2005; Voight et al. 2006; Ferrer-Admetlla et al. 2014; Garud et al. 2015)), tend to be biased towards detecting hard sweeps because their more prominent genomic footprints are easier to detect (Teshima et al. 2006; Garud et al. 2015; Alachiotis and Pavlidis 2018; Weigand and Leese 2018). Even methods that are capable of detecting soft sweeps often lose potentially valuable information by reducing population genomic diversity to a single statistic (Schrider and Kern 2016). This endeavor is further complicated in populations with complex or unknown demographic histories because certain events, like bottlenecks, can mimic selective sweeps (Simonsen et al. 1995; Jensen et al. 2005; Nielsen et al. 2005). In the last few years, the field has made substantial progress in leveraging powerful machine learning algorithms that have been shown to outperform traditional methods (Flagel et al. 2019; Torada et al. 2019; Adrion et al. 2020; Caldas et al. 2022; Hejase et al. 2022; Mo and Siepel 2023; Whitehouse and Schrider 2023).

Machine learning (ML) approaches yield impressive discriminatory power as they either use a combination of several summary statistics as input (Pavlidis et al. 2010; Lin et al. 2011; Ronen et al. 2013; Schrider and Kern 2016; Alachiotis and Pavlidis 2018; Kern and Schrider 2018; Mughal et al. 2020; Arnab et al. 2023) or bypass this step entirely and train neural networks to work directly on genome alignments (Chan et al. 2018; Flagel et al. 2019; Torada et al. 2019; Adrion et al. 2020; Sanchez et al. 2021). Central to ML approaches are simulated training datasets which can be generated using estimated population-specific demographic histories. Doing so allows researchers to better approximate the distributions of patterns of polymorphisms produced by selection and neutrality even under more realistic and complex scenarios, thereby improving sweep-detection accuracy (Pybus Oliveras et al. 2015; Schrider and Kern 2016). However, this also means that the performance of these approaches is fundamentally tied to the demographic models used to generate the simulations because the classifier effectively “learns” the demographic assumptions that are provided. If the demographic history is misspecified, the classifier may misinterpret neutral patterns as selection or fail to identify true sweeps, meaning that accuracy is influenced by the chosen demographic model. There are, however, some methods that remain robust to demographic misspecification by focusing on spatial patterns of variation that signal selective sweeps, rather than relying on local levels of genetic variation (Schrider and Kern 2016). This is because demographic events typically leave signatures on a genome-wide scale, while strong selective sweeps may be expected to produce a localized perturbation in patterns of genetic diversity regardless of the population’s demographic history. Finally, some ML approaches have the added capacity to distinguish between hard and soft sweeps (Kern and Schrider 2018; Mughal and DeGiorgio 2019). This functionality is especially important in large, diverse populations where greater levels of standing variation and/or a higher population-scaled mutation rate may mean that adaptation predominantly occurs through soft sweeps (Messer and Petrov 2013).

As a globally distributed vector species with large population sizes and high levels of genetic diversity, the mosquito *Aedes aeygpti* poses a major threat to public health (Schmidt et al. 2020; Gómez-Palacio et al. 2024; Kent et al. 2025; Lozada-Chávez et al. 2025). *Ae. aegypti* transmits yellow and dengue fever, Zika, and chikungunya, making a thorough understanding of natural selection in this species critical for guiding mitigation and vector control strategies. Due to a combination of climate change which is making regions more suitable for *Ae. aegypti,* increased globalization, and their impressive capacity as an invasive species, *Ae. aegypti* populations are expanding in parts of Europe, Central America, East Africa, the United States, and Canada (Ryan et al. 2019; Iwamura et al. 2020; Kolimenakis et al. 2021; Laporta et al. 2023). In addition to novel environmental stressors experienced during range expansion, *Ae. aegypti* has also encountered strong selective pressure in the form of insecticides (Love et al. 2023; Ware-Gilmore et al. 2023). Much of our understanding of the development of insecticide resistance in *Ae. aegypti* comes from studies in Asia and the Americas, whereas data from Africa remain comparatively limited. In some parts of the Americas, *Ae. aegypti* has experienced insecticide-based vector control for nearly a century while in Africa, *Ae. aegypti* has not been targeted by focused vector control strategies to the same extent as malaria vectors (Weetman et al. 2018; Wilson et al. 2020; Yaméogo et al. 2024). Nonetheless, increasing levels of insecticide resistance have now been documented across both Africa and the Americas, despite substantial variation in the intensity, duration, and documentation of insecticide use in these regions. This, coupled with their population sizes, genetic diversity levels, and capacity to develop resistance to insecticides in as little as five generations, suggests that signatures of positive selection may be prevalent (Martins et al. 2012; Matthews et al. 2018; Thornton et al. 2020; Kent et al. 2025). However, efforts to detect such signals are complicated by the species’ varied and complex demographic histories (Nielsen et al. 2005; Kent et al. 2025).

Originally native to Africa, *Ae. aegypti* has, over the last four centuries, expanded its geographic range to include most of the world’s tropical belt (Ryan et al. 2019; Rose et al. 2023). The movement of *Ae. aegypti* from Africa to the Americas involved multiple introductions, bottlenecks, and expansion events and was likely facilitated by the high ship volume to the Americas during the Transatlantic Slave Trade (Tabachnick 1991; Brown et al. 2014; Powell et al. 2018). Population size estimates and nucleotide diversity in African *Ae. aegypti* population samples are approximately 10^5^–10^6^ and 0.0370, respectively (Rose et al. 2023; Kent et al. 2025). These estimates are generally larger than those in *Drosophila melanogaster*—a species known for its large population sizes and high levels of genetic diversity (Terhorst et al. 2017; Kapopoulou et al. 2018). Most of the genetic studies performed on *Ae. aegypti* have relied on only a small number of genes (Bennett et al. 2016), mitochondrial DNA (Gonçalves da Silva et al. 2012), microsatellites, or reduced representation sequencing (Brown et al. 2014), and only recently have whole genome-based resequencing and exome-based sequencing been performed (Crawford et al. 2017; Matthews et al. 2018; Kelly et al. 2021; Lozada-Chávez et al. 2025). This is largely due to the size (1.3 Gb) and repetitive nature of its genome which poses a significant challenge (Matthews et al. 2018).

Because whole-genome data for *Ae. aegypti* have only recently become available (Rose et al. 2020; Kelly et al. 2021; Love et al. 2023), very little is known about recent positive selection in this species. Currently, sweep scans in *Ae. aegypti* based on single-summary statistical approaches have reported positive selection in genomic regions linked to human preference (Rose et al. 2020), increased tolerance to egg desiccation (Venkataraman et al. 2022), and insecticide resistance (Saavedra-Rodriguez et al. 2019; Love et al. 2023; Schmidt et al. 2024; Lozada-Chávez et al. 2025). Among these, the best-characterized insecticide-resistance genes include the voltage-gated sodium channel (*VGSC*), the acetylcholinesterase gene *ace-1*, and metabolic detoxification genes such as cytochrome P450s and glutathione S-transferases (Fukami 1980; Mutero et al. 1994; Williamson et al. 1996; Daborn et al. 2002). However, given their large population sizes, high levels of genetic diversity, and growing evidence to suggest that IR-increasing alleles may spread via soft sweeps in several insect species (Garud et al. 2015; Xue et al. 2021; Muralidhar and Veller 2022), it is likely that traditional methods may be missing informative signatures of selection. Here, we leverage a robust machine learning approach (Kern and Schrider 2018) to identify genomic targets of positive selection from environmental and anthropogenic stressors in *Ae. aegypti*. We find that we are able to accurately detect both hard and soft sweeps in four globally distributed population samples with complex population size histories. Importantly, we identify novel insecticide resistance (IR) genes which will improve our understanding of the mechanisms of IR evolution and uncover evidence that soft sweeps may play a central role in adaptation in *Ae. aegypti*.

## RESULTS

### Accurate detection of hard and soft selective sweeps in *Aedes aegypti*

To identify genomic targets of selective sweeps in the yellow-fever mosquito, *Aedes aegypti*, we used a machine learning approach, diploS/HIC (Kern and Schrider 2018), which seeks to discriminate between hard sweeps, soft sweeps, regions linked to hard or soft sweeps, and purely neutrally evolving regions. We used previously published genomic data from Love et al., (2023) that examined mosquito population samples from Brazil (Santarém), Colombia (Río Claro and Cali), Gabon (Franceville), Senegal (Ngoye), and Kenya (Kaya Bomu; Figure 1). For each population sample, we applied a diploS/HIC classifier that was trained on simulations following the population size trajectory estimates from Kent et al. (2025) (Supplemental Figure 1) that were estimated using SMC++ (Terhorst et al. 2017). SMC++ has been shown to infer accurate population size histories across broad time frames by combining the use of the SMC framework, which is most accurate in the deep past, with allele frequency information from a larger sample, which can improve estimation accuracy in the relatively recent past (Liu and Fu 2015; Patton et al. 2019; Liu and Fu 2020). Kent et al. further validated their *N_e_* findings by running Stairway Plot 2 (Liu and Fu 2020) which showed similar patterns to those generated through SMC++. A generation mutation rate of 4.85 × 10^−9^ and a generation time of 0.067 years was used according to Rose et al. (2020). Our training simulations also included variation in mutation rates, recombination rates, and selective parameters such as the selection coefficient and the timing of the sweep (see Methods). We specifically chose parameters designed to model recent and strong selective sweeps capable of substantially altering patterns of diversity.

**Figure 1.**
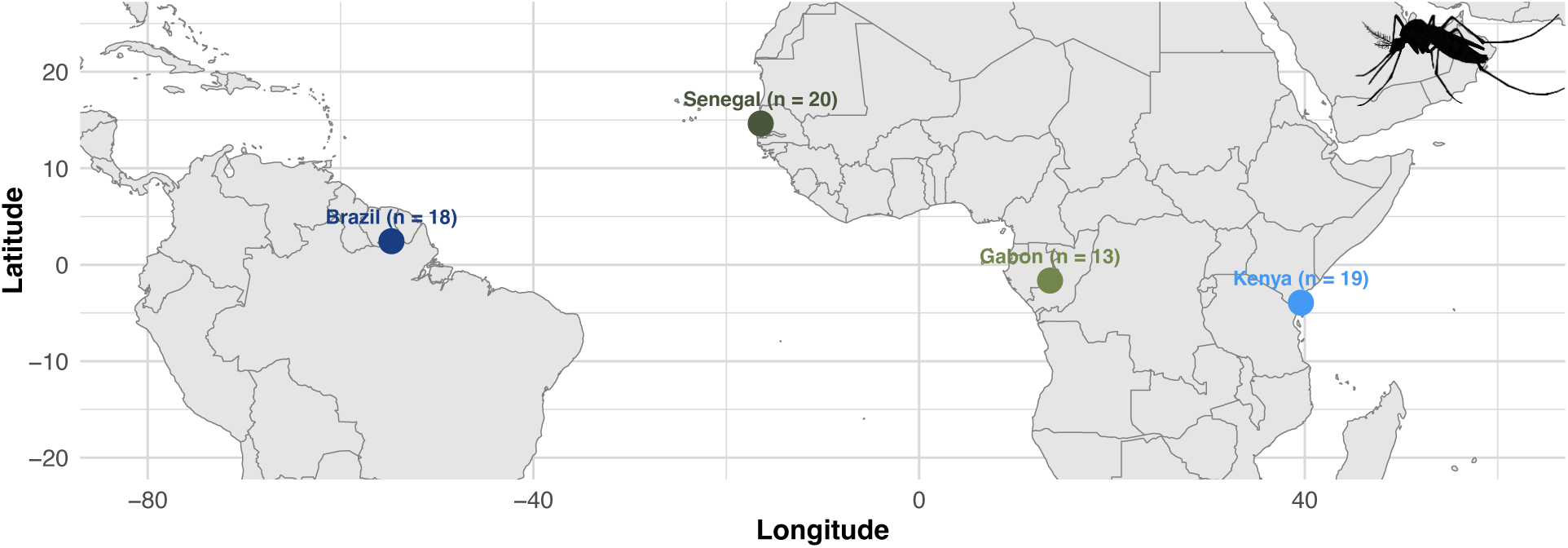
Map of sampling locations including the number of samples analyzed from each site. We have omitted the two sampling sites from Colombia as they were not included in the final analysis.

We assessed the performance of our classifiers through Receiver Operating Characteristic curves (ROC), Precision Recall (PR) curves, and confusion matrices (Figure 2 and Supplemental Figure 2). ROC curves measure true positive versus false positive rates across classification thresholds. An ideal classifier has a ROC curve that resembles a step-function with a true positive rate (or recall) as close to 1.0 as possible while achieving a false positive rate as close to 0 as possible. A PR curve measures the precision (or positive predictive value) against recall, with a good classifier being able to maintain high precision even at classification thresholds that yield high rates of recall. Here, our curves evaluate our models’ performance on the binary task of distinguishing selective sweeps (whether hard or soft) from unselected regions (whether sweep-linked or fully neutrally evolving). Classifier accuracy was evaluated using held-out simulated test datasets generated under the same prior parameter distributions used for training, meaning that the reported metrics reflect performance on data drawn from the same underlying assumptions as the training set. The excellent quality of our classifiers is clearly discernible by the ROC and PR curves which respectively show an area under the curve (AUC) value greater or equal to 0.97 and an Average Precision (AP) value greater or equal to 0.95 in all population samples except for the two Colombia population samples. The area under the ROC curve quantifies the classifier’s ability to discriminate between classes across all thresholds, whereas the average precision summarizes performance across all recall values (equal to the area under the precision-recall curve).

**Figure 2.**
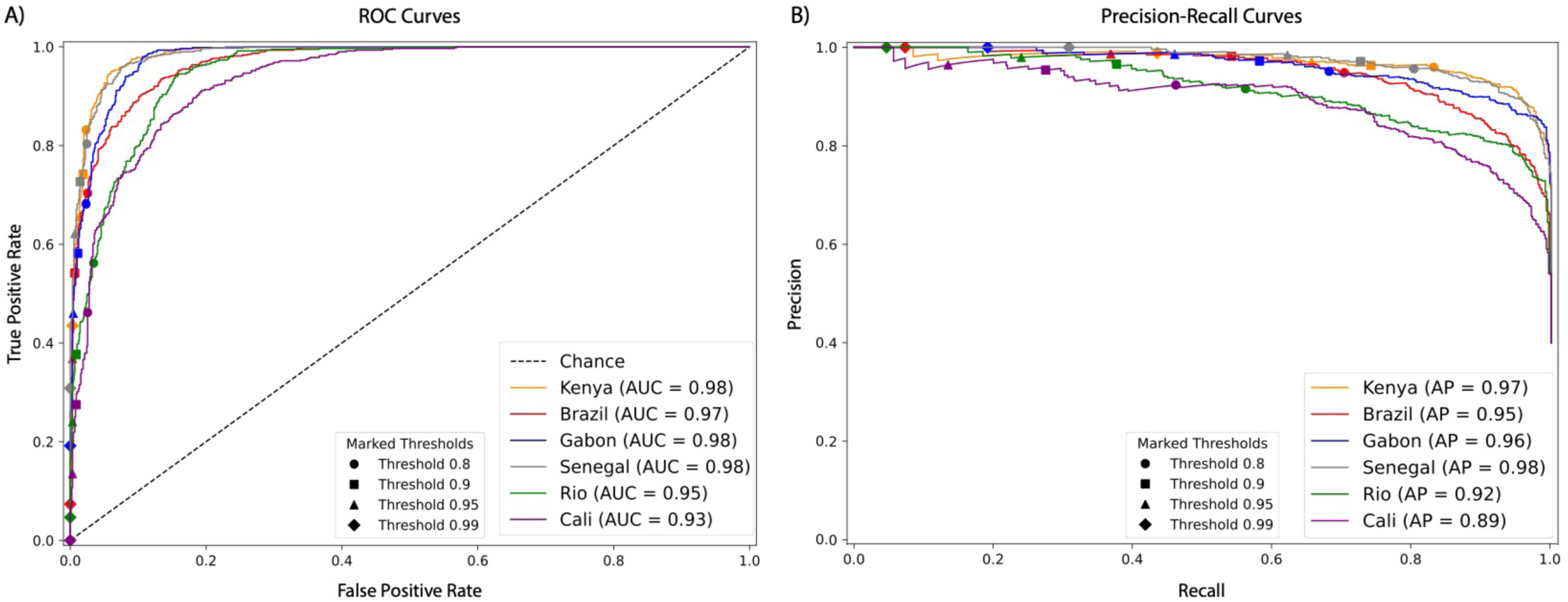
ROC curves and precision-recall curves summarizing the performance of each population samples’ classifier. A) ROC curves showing the true and false positive rates for the binary classification task of distinguishing between selective sweeps (hard and soft) vs unselected regions (sweep-linked and neutral) with varying threshold cutoffs highlighted with different shapes. B) Precision-recall curves showing the classifiers performance at the same task (sweep vs unselected) with varying threshold cutoffs highlighted with different shapes. The precision is defined by the fraction of regions classified as sweeps that truly were sweeps and the recall is defined by the true positive rate. For the Cali classifier, there were no windows classified as a sweep with probability ≥0.99, so no marker is included on Cali’s precision-recall curve for that threshold.

While ROC and PR curves are useful for summarizing the overall accuracy of a binary classifier, for multi-class inferential models like our trained diploS/HIC neural networks, these curves are limited because they collapse diploS/HIC’s five classes down to two (selected and unselected regions), and thus they are not informative about the precise type of errors made within these two meta-classes. For example, if a hard sweep is correctly detected, how often is it misclassified as soft? Or, when neutrally evolving regions are misclassified as sweeps, are they generally classified as hard sweeps or soft sweeps? To address these questions, we calculated confusion matrices, which show the fraction of test examples from each of the five classes that are correctly assigned to a given class. However, unlike the ROC and PR curves, the confusion matrices only show results from a single class membership probability threshold. We therefore made confusion matrices for a range of thresholds for each population sample (see Methods). Note that the class membership probabilities, which are produced by the softmax activation function from the final layer of the diploS/HIC neural network, may not necessarily be well calibrated such that 80% of examples classified as a hard sweep with 80% probability will be true hard sweeps.

In each population sample our ability to distinguish simulated sweeps from neutral variation was strong and improved further when imposing sweep probability thresholds (e.g., < 7.5% of neutral windows were misclassified as sweeping in each population sample with no sweep probability cutoff versus < 3.1% in each population sample at the 0.80 cutoff; Supplemental Figure 3). When neutrally evolving regions were misclassified, they tended to be misclassified as soft or soft-linked, with a combined misclassification rate < 7.04% at the 0.80 cutoff across all population samples except the Cali and Río Claro samples from Colombia, which showed lower accuracy (see below). Neutrally evolving regions were never classified as hard sweeps at the 0.80 cutoff in any population other than Río Claro, and the rate at which they were misclassified as soft sweeps (in all populations except for those from Colombia) was < 1.7%. Our diploS/HIC classifiers also demonstrated adequate power to discriminate between hard and soft sweeps. When sweep regions were misclassified, they tended to be misclassified as the reciprocal sweep condition (i.e., soft sweeps were misclassified as hard sweeps and vice versa). This same trend occurs in sweep-linked regions. These results indicate that our classifiers were able to robustly distinguish sweeping regions from neutrally evolving or sweep-linked regions.

Overall, the accuracy of our classifiers was higher in African population samples than the South American population samples (Supplemental Figures 2-6). For example, in Kenya with no sweep probability cutoff, diploS/HIC correctly classified 88% of hard sweeps and 68% of soft sweeps (Supplemental Figure 2); with a 0.80 threshold imposed this increased to 88% and 72%, respectively. This improvement is observed because simulations classified as sweeps, but with a combined sweep probability lower than the classification threshold, are treated as uncertain and are thus omitted from the calculation, and those high-confidence predictions that remain are more likely to be accurate. When applying cutoff values of 0.90, 0.95, and 0.99 to the Kenya classifier, accuracy values for hard sweeps stayed relatively constant at 88%, 88%, and 90%, respectively, as did soft sweeps at 71%, 69%, 66%. Conversely, for Cali (Supplemental Figure 2) with no cutoff, diploS/HIC correctly classified 80% of hard sweeps and 33% of soft sweeps and increasing the threshold to more stringent values did not improve these values (e.g., at 0.90, 57% of hard sweeps and 25% of soft sweeps were recovered). At the highest threshold of 0.99, there were no hard or soft sweep simulations for Cali passing the threshold, so the fraction of sweeps correctly predicted could not be calculated. Performance for the other Colombian sample (Río Claro) was similarly poor (Supplemental Figure 2-6). The poorer classifier performance obtained on simulated test data for the Colombian populations samples was expected due to their demographic histories which included stronger and more recent bottlenecks than the other population samples (Supplemental Figure 1). These bottlenecks reduce genetic diversity and distort genealogies in a manner that can resemble positive selection, making sweep detection inherently more difficult (Jensen et al. 2005; Schrider and Kern 2016). Given the performance of Colombia’s classifiers, we chose to focus our subsequent analyses on the other four population samples (Brazil, Gabon, Kenya, and Senegal), for which our simulated test data suggested that we could detect and classify sweeps with sufficient accuracy.

### Soft sweeps appear to predominate over hard sweeps in *Aedes aegypti*

We classified a total of 4598, 4599, 4599, and 4599 windows that passed our data filtering cutoffs in the Brazil, Gabon, Senegal, and Kenya population samples, respectively. Of these windows, 165 were classified as sweeps in Brazil, compared to 150 in Gabon, 79 in Senegal, and 363 in Kenya (Table 1 and Figure 3A). To construct higher-confidence sets of sweep candidates, we then imposed increasing posterior probability cutoffs on these windows (Table 1). For example, when using a cutoff of 0.95, there were 24 sweep windows in Brazil, 18 in Gabon, 17 in Senegal, and 65 in Kenya. These windows comprise a set of 106 distinct candidate sweeps, some of which were shared by more than one population sample: 1 (0.94%) was shared across all population samples, 1 (0.94%) was shared among the African population samples, and 93 (87.7%) were population-specific sweeps (Figure 3, Supplemental Figure 7). The remaining 11 (10.4%) sweeps were present in more than one population but did not fit into the categories above.

**Figure 3.**
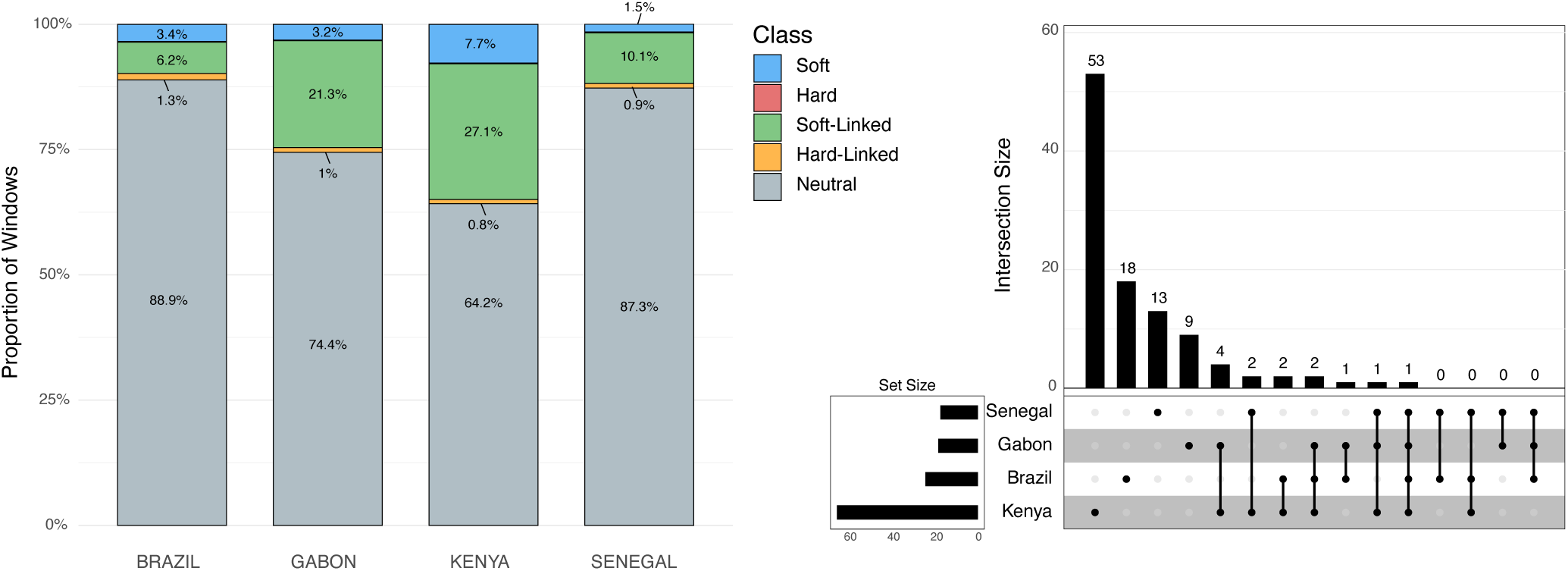
(Left) The proportions of each classification type per population sample with no threshold cutoff applied. diploS/HIC predicted 4,088 neutral, 286 soft-linked, 59 hard-linked, 157 soft, and 8 hard windows for Brazil. For Gabon, predictions included 3,423 neutral, 981 soft-linked, 45 hard-linked, 145 soft, and 5 hard windows. For Kenya, predictions included 2,952 neutral, 1,245 soft-linked, 39 hard-linked, 355 soft, and 8 hard windows. For Senegal, diploS/HIC predicted 4,013 neutral, 464 soft-linked, 43 hard-linked, 71 soft, and 8 hard windows. (Right) Upset plot showing the intersection of selective sweep locations across the four population samples in this study at a posterior probability cutoff of 0.95.

**Table 1.**
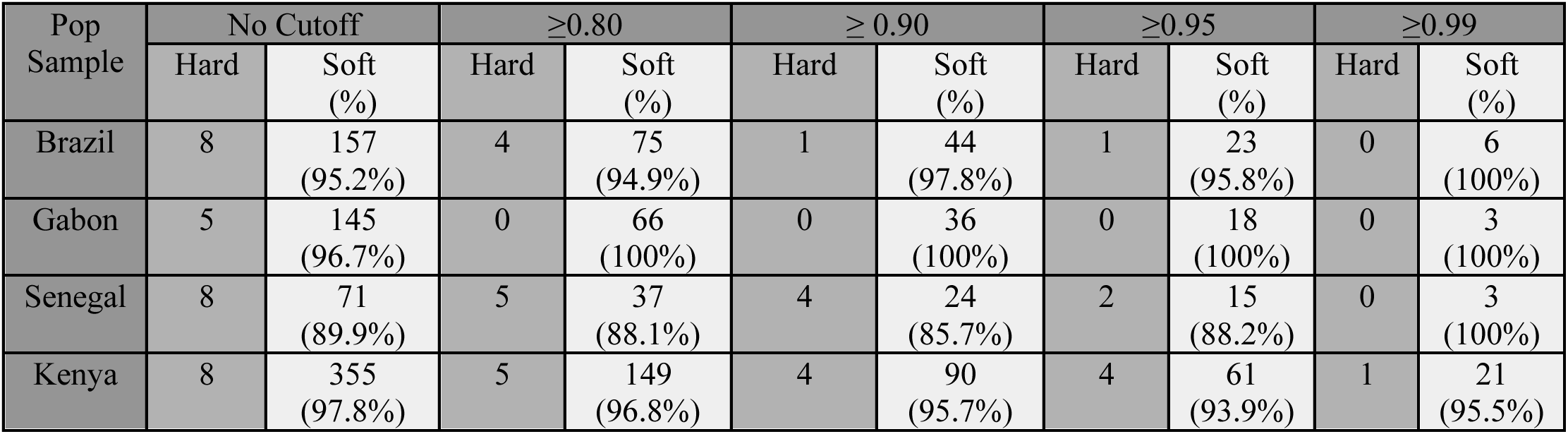
The number of hard and soft sweeps per population sample at different posterior probability thresholds.

Across all population samples, the number of soft sweeps was substantially higher than the number of hard sweeps and this pattern did not change with more stringent posterior probability thresholds (Table 1 and Figure 3). We further evaluated the robustness of this finding by applying a conservative approach using the error rates in our confusion matrices to estimate the proportion of sweeps that were soft after accounting for hard sweeps that may have been misclassified as soft, and to estimate potential false positive soft sweeps that are truly neutrally evolving—this approach may underestimate the fraction of soft sweeps because we ignore the possibility of soft sweeps being misclassified as hard (see Methods). Even after accounting for these estimated error rates, we found that soft sweeps predominate at every threshold and that the relative proportion of soft sweeps generally increases or remains stable with more stringent threshold cutoffs. Indeed, at every threshold and every population sample, more than 72% of all sweeps are predicted to be soft (Supplemental Table 1). In all population samples other than Senegal, this (conservative) estimate was even higher: > 83.5% across all thresholds.

To further measure the degree of confidence in our sweep calls, we calculated *q*-values, a false discovery rate (FDR)-adjusted *p*-value derived from procedure that is more powerful than the Benjamini–Hochberg FDR method (Storey 2002). We did this for each sweep window based on the combined posterior probabilities of hard and soft sweeps and our predicted false positive rates obtained from simulations for each population sample (Methods; see Supplemental Table 2 for the sweep candidates and their associated *q*-values). Notably, for those sweep windows with a combined posterior probability of ≥ 0.95, our *q-*value estimate could not be distinguished from zero (i.e., our neutral test simulations never achieved a sweep probability above this threshold) in all population samples except for Kenya (where sweeps passing the 0.95 threshold had an estimated *q*-value of ∼0.4).

### Insecticide Resistance Genes as Likely Targets of Selection

When examining candidate selective sweeps, we separately considered two sets of regions: shared sweeps (those with a posterior sweep probability of ≥ 0.95 in more than one population sample) and population-specific sweeps (those where only one population sample had a sweep probability ≥ 0.95). These two sets contained 13 and 93 sweep windows, respectively. Many of these candidate regions include genes with functions potentially linked to insecticide resistance (IR). To better interpret their relevance, we categorized these genes into two groups: **well-characterized IR genes** and **putative or emerging IR candidates**. We assigned genes to these groups based on the nature and extent of evidence for their relevance to IR in the literature, including: (1) functional similarity to known IR genes, (2) differential expression in response to insecticide exposure, and (3) signatures of selection reported in other insect species under insecticide pressure. It is important to note that while we can identify compelling candidate genes within these sweep regions, there are often multiple genes in and around these windows, and we thus cannot conclude with certainty which gene is the target of selection.

#### Shared Sweep Windows Containing Putative or Emerging Insecticide Resistance Candidate Genes

Of the 13 windows that were shared between two to four population samples, three windows contain candidate *Ae. aegypti* insecticide-resistance genes (see Supplemental Table 3 for the complete list). First, we found a high-confidence soft sweep shared in Kenya and Gabon (chr2: 452,250,001-452,500,000) that contains an ankyrin repeat domain-containing protein 29. This same region was predicted to be a soft sweep in Brazil but its sweep probability of 0.94 did not meet our threshold of 0.95. The ankyrin protein domain is a common protein-protein interaction motif that is present in numerous structural proteins and is involved in cytoskeletal anchoring and mechanosensation. Importantly, ankyrin proteins have been shown to directly interact with the well-established target for insecticides, the voltage-gated sodium channel VGSC (Williamson et al. 1996). In a study on pyrethroid resistant *Ae. aegypti* from Mexico, genes containing the ankyrin-domain were highly associated with pyrethroid resistance (Campbell et al. 2019). Similarly, in pyrethroid-resistant populations of the mosquito *Anopheles funestus* from Senegal, an ankyrin repeat domain protein was one of the most highly overexpressed genes (Samb et al. 2016). In *Ae. aegypti*, an ankyrin repeat domain-containing protein was found to be associated with pyrethroid resistance (Cosme et al. 2022), and signatures of selection in two regions containing ankyrin genes (along with other IR candidates) was also reported by Love et al., (2023). Similar patterns have also been observed in other insect species (Kwiatkowska et al. 2013; Gouesbet et al. 2025), further highlighting the role of these gene families in pyrethroid resistance.

The other two shared sweep windows containing putative IR genes were also classified as sweeps in Kenya and Gabon. The first contained a gene that encodes for the RNA binding protein Split ends (*spen;* chr2: 143,500,001-143,750,000) which provides a protective role against cytotoxicity from the herbicide paraquat in *Drosophila* (Girard et al. 2020; Bresgen et al. 2023). Although paraquat is an herbicide, it is known to cause oxidative stress in the mosquito *An. gambiae* (Champion and Xu 2018; Tarimo et al. 2018). The second window contains *thioredoxin-2* (chr2: 45,000,001-45,250,000), a key mitochondrial protein that regulates cellular redox and is protective against oxidative stress, a common consequence of insecticide exposure, in *Drosophila* (J. Svensson and Larsson 2007). Interestingly, the thioredoxin system recycles oxidized glutathione to reduced glutathione, which can then be used by the well-established IR genes glutathione S-transferases (GSTs) for detoxification (Tarimo et al. 2018). Other components of the thioredoxin system, like thioredoxin peroxidase, have been shown to protect insects from insecticide-induced oxidative injury (Zhao et al. 2022; Gao et al. 2025) and *thioredoxin* is listed as of the candidate IR genes on the *Anopheles gambiae* 1000 Genomes Project Selection Atlas (Clarkson et al. 2020).

#### Population-Specific Sweep Windows Containing Well-Characterized Insecticide Resistance Genes

We classified 93 windows as a sweep with ≥ 95% probability in a single population sample. Of these 93 windows, six contain genes with well-characterized IR-related functions. We describe three of these genes below (See Supplemental Text for descriptions of the other three genes) and explore their role in insecticide resistance.

One window classified as a soft sweep in Senegal, located at chr1:271,250,001-271,500,000, contains three cytochrome P450 genes: *Cyp 6a8, Cyp 6a13, Cyp 6a14* (Figure 4). Cytochrome P450 monooxygenases (CYPs) are critical for metabolic detoxification and have been implicated in insecticide resistance in many species, including mosquitoes (Balabanidou et al. 2016; Rahman et al. 2021; Yang et al. 2021). *Cyp6a8* specifically has been shown to be upregulated in response to *Piper nigrum* and DDT in *D. melanogaster* (Maitra et al. 1996; Jensen et al. 2006). Notably, this sweeping window also contained several SweepFinder Composite Likelihood Ratio (CLR) peaks (Nielsen 2005), with the highest value > 750. We also detected a soft sweep in Kenya in a window that contains *cytochrome b5 reductase 4* (chr2:156,500,001-156,750,000). Cytochrome P450’s require cytochrome b5 reductase to function as an electron-transfer intermediate and the latter is therefore likely to be involved in IR (Zhao et al. 2012). Moreover, it has been shown to be upregulated in response to phenobarbital in the cotton bollworm, *Helicoverpa armigera* (Zhao et al. 2012) and has been linked to cyantraniliprole resistance in the tomato pest *Tuta absoluta* (Ullah et al. 2025).

**Figure 4.**
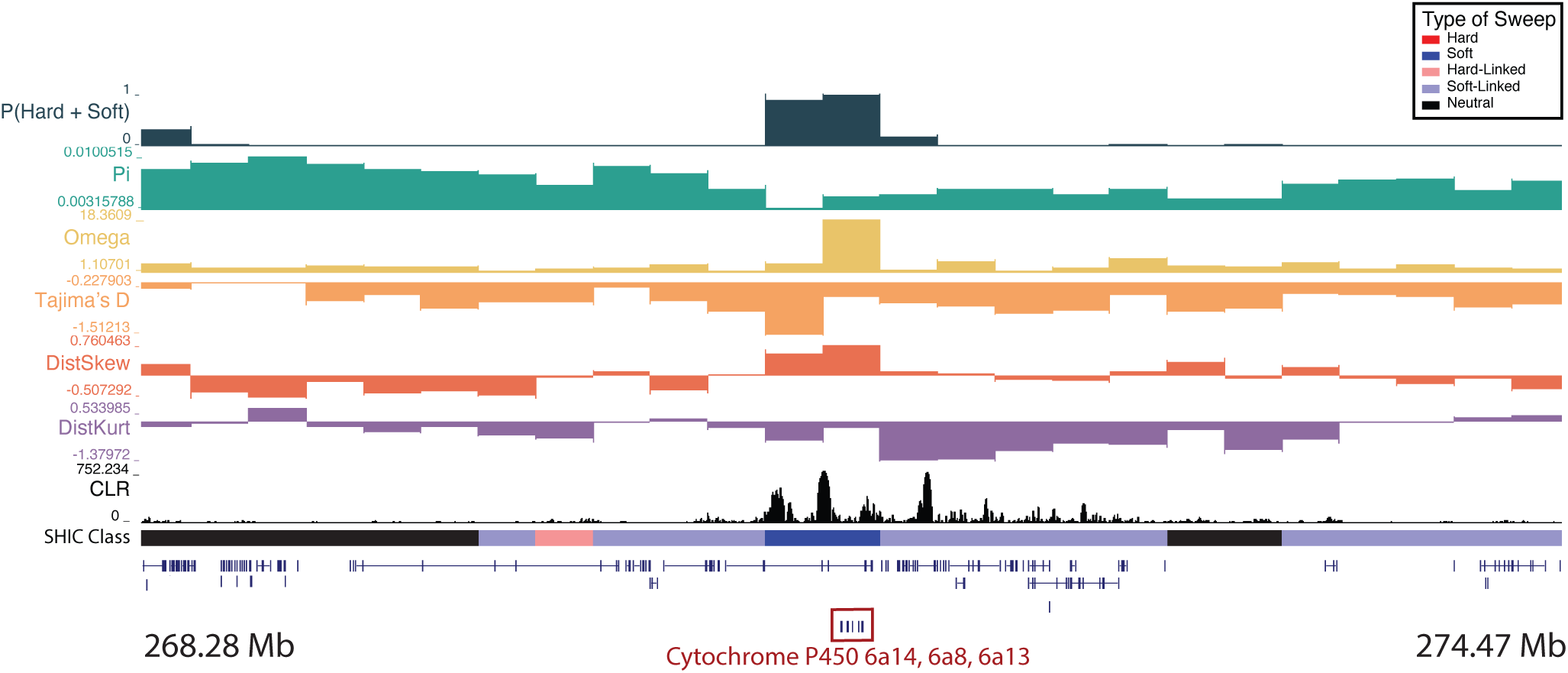
A soft sweep in Senegal at three cytochrome P450 genes: *Cyp6a14*, *Cyp6a8*, and *Cyp6a13* on chromosome one. The diploS/HIC classification track shows the class with the highest posterior probability, with soft sweeps as dark blue, soft-linked regions as light blue, hard sweeps as red, hard-linked as light red, and neutrally evolving regions in black. Above the diploS/HIC classifications are a subset of the summary statistics used by the classifier.

We detected a soft sweep in Brazil in a window containing glutathione S-transferase (chr2:446,250,001-446,500,000), *GSTD7*, and that had a corresponding CLR peak with values > 460 (Supplemental Figure 8). Glutathione S-transferase epsilon clusters (GSTe) are another group of metabolic enzymes that are well-known for their role in resistance to several classes of insecticides (Enayati et al. 2005), and other *GSTe* genes have previously been found to exhibit strong signatures of positive selection (Love et al. 2023; Schmidt et al. 2024). *GSTD7*, specifically, was found to be under selection in the fall armyworm, *Spodoptera frugiperda* (Tessnow et al. 2025). Moreover, knockdown of GSTD7 resulted in higher mortality post-imidacloprid exposure in the silver whitefly *Bemisia tabaci* (He et al. 2018), and *GSTD7* is highly expressed in *Drosophila suzukii* after exposure to malathion (Hamby et al. 2013).

Because we also wanted to explore two of the most well-known IR genes, our results next focus on two major selective sweeps previously reported in Schmidt et al. (2024), spanning the voltage-gated sodium channel gene (*VSSC/VGSC*) and a cluster of 15 glutathione S-transferase genes (*GST*)—regions likewise identified as sweeps in Love et al. (2023). Genome-browser visualizations of both regions are provided in Supplemental Figures 9 and 10. In Brazil, we detected a sweep signal in the *VGSC* region, although its combined posterior probability (0.91) fell slightly below our predefined 0.95 threshold. The magnitude of this sweep, visible as a multi-megabase Tajima’s *D* valley, suggests that a larger diploS/HIC window size would be required for more confident classification. As this sweep is easily detectable without the aid of a machine-learning framework incorporating many summary statistics, we deemed it justified to employ a smaller window size so as to prioritize the detection of less expansive sweeps. We likewise detected a sweep signal in the *GST* gene cluster in Brazil, with one diploS/HIC window surpassing our threshold (posterior probability = 0.96). Although this sweep is also quite large, its signature is not as broad as that of *VGSC*, so our method was able to examine the entire sweep region and detect it with higher confidence.

#### Population-Specific Sweep Windows with Putative or Emerging Insecticide Resistance Candidate Genes

Of these 93 windows that were classified as high-confidence sweeps, 13 contain genes with putative or emerging IR-related functions. We describe seven of these genes below and explore their role in insecticide resistance (see Supplemental Table 4 for the complete list). The remaining six genes and associated role in IR are included in the Supplementary Text.

We found three high-confidence windows that were classified as soft sweeps in Kenya which contained *arrestin* (chr2:246,250,001-246,500,000; Supplemental Figure 11) and overlapped with the *sodium/potassium/calcium exchanger Nckx30C* (chr2:288,750,001-289,000,000; Figure 5). In vertebrates, arrestins regulate the signaling and trafficking of G-protein-coupled receptors (Gurevich and Gurevich 2006), and in the mosquito *Culex pipiens pallens*, *arrestin* expression was higher in deltamethrin-resistant strains and knockdown via siRNA resulted in decreased viability after deltamethrin treatment (Sun et al. 2012). *Nckx30C* is likely involved in the maintenance of calcium homeostasis (Haug-Collet et al. 1999; Webel et al. 2002) and was found to be differentially regulated in insecticide resistant strains of *An. gambiae* (Vontas et al. 2005). The last window classified as a hard sweep in Kenya and contains *mitochondrial protoheme IX farnesyltransferase* (chr2:262,250,001-262,500,000), or *COX10*. *COX10* encodes for an enzyme that catalyzes the farnesylation of heme, which plays a crucial role in cytochrome c oxidase (COX) function (Diaz et al. 2006). COX genes have been linked to insecticide resistance in a number of species (Steele et al. 2018), including *D. melanogaster* (Song and Scharf 2009), the housefly *Musca domestica* (Sacktor 1951), the German cockroach *Blattella germanica* (Pridgeon and Liu 2003), and *Ae. aegypti* (Pridgeon et al. 2009).

**Figure 5.**
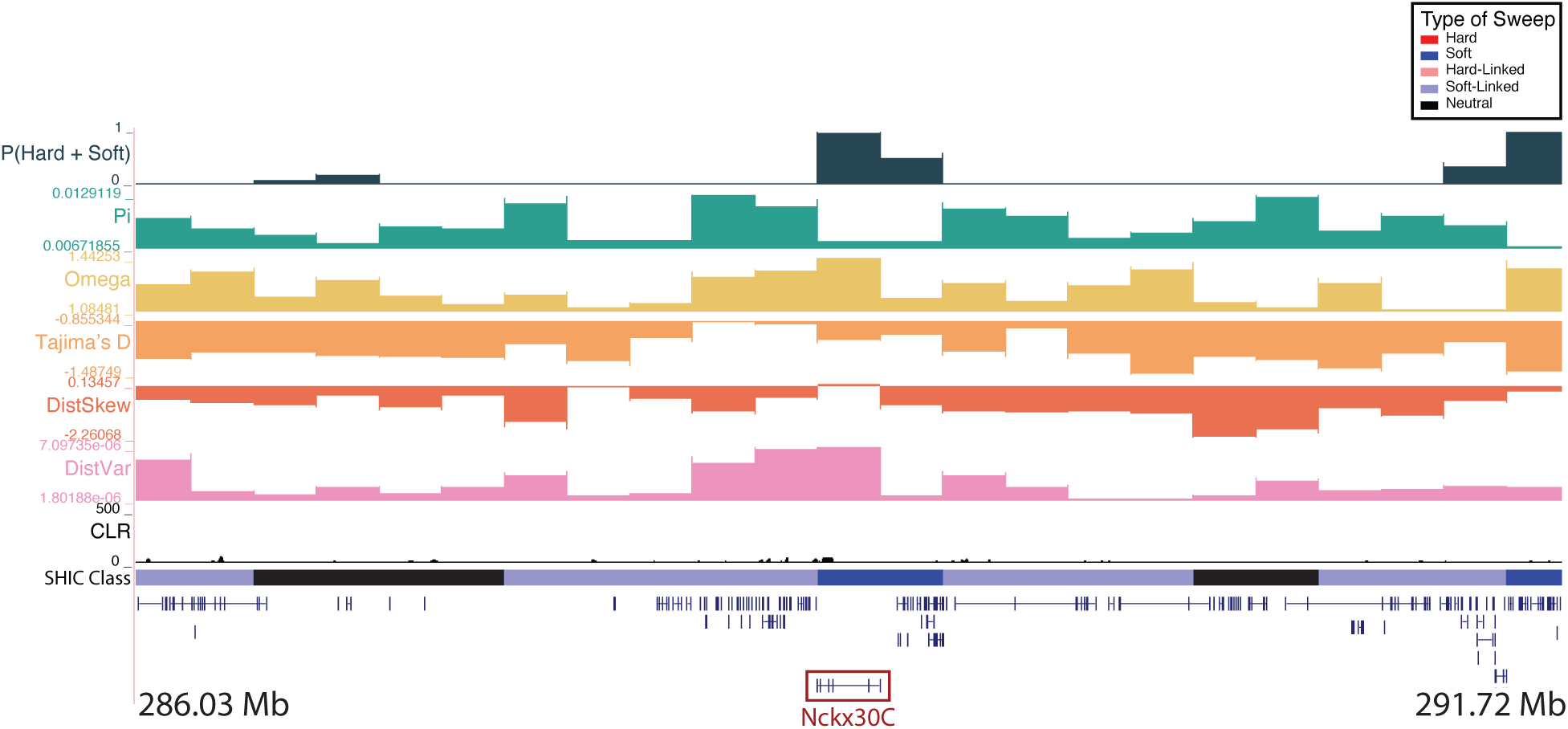
A soft sweep in Kenya at the *sodium/potassium/calcium exchanger Nckx30C* on chromosome two. The diploS/HIC classification track shows the class with the highest posterior probability, with soft sweeps as dark blue, soft-linked regions as light blue, hard sweeps as red, hard-linked as light red, and neutrally evolving regions in black. Above the diploS/HIC classifications are a subset of the summary statistics used by the classifier.

In Brazil, we found sweeping windows that contain or overlap the *muscle calcium channel subunit alpha-1* (chr2:225,750,001-226,000,000; Supplemental Figure 12), *potassium voltage-gated channel subfamily KQT member 1* (chr2:144,750,001-145,000,000), *neuroligin-1* (chr3:351,750,001-352,000,000), and *thioredoxin-related transmembrane protein 1* (chr3:12,250,001-12,500,000). Of these four windows, three are classified as soft sweeps while the first window is classified as a hard sweep. Further, the second, third, and fourth window all contain CLR peaks with values greater than 475, 280, and 900, respectively. The first window was also initially classified as a soft sweep in Kenya, although the posterior probability (0.83) did not meet our threshold. Although the role of both the *muscle calcium channel subunit alpha-1* and the *potassium voltage-channel subfamily KQT member 1* in IR is not well understood, both mechanosusceptible and voltage-gated channels are confirmed targets of various insecticide and pesticide classes. *Neuroligin-1* is a synaptic cell-adhesion molecule (Song et al. 1999) that is highly associated with pyrethroid resistance in *Ae. aegypti* (Campbell et al. 2019), likely a result of insecticides disrupting excitatory synapses. Similar to the *thioredoxin-2* sweep window described above, *thioredoxin-related transmembrane protein 1* (Trx1) also plays a role in protection against oxidative stress in insects (Zhang et al. 2015) and has been shown to be involved in antioxidant defense in *Apis cerana cerana* (Yao et al. 2014).

We discuss additional candidate sweep regions containing the following putative IR genes: *acyl-CoA synthetase family member 4*, *fatty acyl-CoA reductase 1, membrane-associated progesterone receptor component 1, D(2) dopamine receptor A, transcription factor grauzone,* and *NADH dehydrogenase 1 beta subcomplex subunit 5* in the Supplemental Text.

### No Evidence of Enrichment of Insecticide Resistance Genes or GO Terms in Sweep Windows

To test whether genes previously implicated in insecticide resistance were overrepresented in our high-confidence sweep windows (posterior probability cutoff ≥ 0.95), we used a list of 157 *Ae. aegypti* genes previously implicated in insecticide resistance (Supplemental Table 5). Importantly, this list was generated prior to identifying and annotating candidate sweep windows with diploS/HIC so that the enrichment results would provide an unbiased assessment of whether known IR loci disproportionately overlapped our inferred sweep regions. Across all four populations, we found no statistically significant evidence of known IR genes among sweep regions. In Gabon and Kenya, the overlap between sweep windows and IR genes was equal to or below that expected by chance. In Brazil and Senegal, enrichment was somewhat elevated by not statistically significant (Supplemental Table 6). We also tested for significantly enriched GO terms within our sweep windows (posterior probability cutoff ≥ 0.80). Although several GO categories showed nominally significant enrichment (*P*-value < 0.05), none of these categories survived FDR correction (*q* > 0.05 in all cases; Supplemental Table 7).

## DISCUSSION

Natural selection leaves conspicuous footprints in the genome, and our ability to accurately detect these signatures has significant implications for public health, agriculture, and conservation. While early methods focused heavily on detecting hard selective sweeps, growing evidence supports the widespread occurrence of soft sweeps, particularly in large, diverse populations. Soft sweeps leave a more subtle signature than hard sweeps and are therefore more difficult to detect with traditional approaches (Wilson et al. 2014). Machine learning has emerged as a powerful tool in this space, well-suited to detecting high-dimensional signals of selection and capable of accounting for complex demographic histories that confound inferences. This is an important advancement for species like *Aedes aegypti*—a major disease vector with a global impact on public health—which poses a significant challenge to evolutionary analyses due to its high degree of genetic variation and complex demographic history. Here we leverage the machine learning tool diploS/HIC (Kern and Schrider 2018) and recently published demographic history estimates (Kent et al. 2025) to uncover the targets of recent positive selection in *Ae. aegypti* and provide evidence that soft sweeps may play a central role in adaptive evolution in this important vector species.

This work contributes to a growing body of literature that challenges the traditional view that selective sweeps mainly act on a single-origin *de novo* mutation. Indeed, our work mirrors the findings of similar studies performed on humans (Schrider and Kern 2017), *D. melanogaster* (Garud et al. 2015), *Anopheles gambiae* (Weedall et al. 2020; Xue et al. 2021), and HIV (Feder et al. 2016), that implicate soft selective sweeps as the dominant mode of adaptation (although these studies have proved controversial; see (Harris et al. 2018; Schrider and Kern 2018; Feder et al. 2021; Garud et al. 2021; Johri et al. 2022)). In fact, in all population samples, and at every sweep probability threshold, the percentage of sweeps that were soft was always > 85%. When we applied a conservative approach designed to account for the potential impact of false positives and of soft sweeps being misclassified as hard (while ignoring the possibility of misclassification in the opposite direction), the percentage of soft sweeps was still > 72% in all population samples and thresholds. This suggests that selection in *Ae. aegypti* can act on standing genetic variation instead of waiting for novel mutations to arise, and *Ae. aegypti* populations are therefore able to rapidly respond to new selective pressures like insecticides. Soft sweeps even predominated in Brazil—a population sample that has experienced a protracted bottleneck associated with the introduction from Africa to the Americas (Rose et al. 2023; Kent et al. 2025). This suggests that even bottlenecked populations of *Ae. aegypti* may have sufficient standing variation (or a high enough population-scaled mutation rate; see below) to facilitate rapid adaptation. Out of every population sample, Kenya had substantially more high-confidence soft selective sweeps predicted than any other population sample (61 in Kenya, compared to 23, 18, 15 in Brazil, Gabon and Senegal, respectively, when considering sweeps with a posterior probability ≥ 0.95). This is in line with the findings presented in Kent et al. (2025) where Kenya was inferred to have a larger effective population size than the other population samples and therefore would be expected to experience efficacious natural selection and more soft sweeps.

Our classification approach modeled soft sweeps as occurring via selection on standing variation, which is plausible given the high levels of genetic diversity observed. Alternatively, selection on recurrent *de novo* mutations can also produce sweep signatures similar to that of a soft sweep on standing variation—albeit typically less pronounced–with multiple haplotypes carrying the beneficial allele (Pennings and Hermisson 2006b). Such sweeps are especially likely in species where the number of new mutations entering the population each generation is large, either due to large population size and/or a high spontaneous mutation rate (Pennings and Hermisson 2006b; Karasov et al. 2010; Garud et al. 2015). Given that soft sweeps on recurrent mutations can look very similar to soft sweeps on standing variation in terms of the values of numerous summary statistics (Schrider et al. 2015), our machine learning models may detect signals of both selection on standing variation and on recurrent mutations. However, both of these sweeps are indicative of a population that need not wait for long periods for a *de novo* beneficial mutation to arise following a change in the selective environment.

On the other hand, there are several phenomena that may result in signatures of soft sweeps even in the absence of rapid adaptation. First, patterns of genetic diversity in a genomic region flanking a hard sweep are expected to mirror that of a soft sweep (the “soft shoulder” effect; (Schrider et al. 2015)). However, this is an unlikely explanation for our results given that we detected very few hard sweeps in our population samples, and diploS/HIC is designed to account for this shoulder effect by examining polymorphisms across a larger window containing the sweep (Schrider and Kern 2016). Second, allelic gene conversion can “soften” hard selective sweeps by transferring the beneficial mutation onto multiple haplotypes (Jones and Wakeley 2008; Schrider et al. 2015; Schrider 2023). This is more common in large populations where there is more time for gene conversion events to occur (Schrider 2023). Third, the dearth of hard sweeps across all population samples may result from the inherent difficulty in accounting for spatial population structure when detecting selective sweeps. In low-dispersal scenarios (which are common in species with wide geographic ranges) an adaptive mutation cannot rise in frequency as rapidly as it would in a panmictic population. As a result, hard selective sweeps can become enriched in intermediate-frequency variants which can cause them to resemble soft sweeps (Chotai et al. 2024). Thus, the expectation for large populations like those examined here might be that we would detect more soft sweeps than hard sweeps even without selection acting on standing genetic variation or recurrent mutations. Finally, we note that demographic model misspecification can diminish accuracy in discriminating between hard and soft selective sweeps. To mitigate this effect, we have trained our classifiers using demographic models estimated from the same population samples examined here, and for which two different demographic estimation methods yielded qualitatively similar results (Kent et al. 2025). Nonetheless, there is direct evidence that *Ae. aegypti* can adapt rapidly to insecticides and Martins et al. (2012) and Lozada-Chávez et al. (2025) recently reported evidence of selection in out-of-Africa population samples on variants that are segregating in Africa. Thus, true soft sweeps may therefore represent the most parsimonious explanation for the patterns we observe, and at least some of these may be due to selection on standing genetic variation.

While the interpretation of soft sweep signatures is not straightforward, the above arguments suggest that it is essential for selection scans in large/diverse populations to consider soft sweeps in order to detect a larger fraction of the targets of recent positive selection. Previous studies have detected selection on several IR-related genes, including the voltage-gated sodium channel gene (*VGSC*), glutathione S-transferases (GSTs), *ace-1,* carboxylesterases, and many cytochrome P450 genes (Love et al. 2023; Schmidt et al. 2024). While these studies have provided insight into how mosquitoes develop resistance to insecticides (e.g., through metabolic detoxification or variation in the insecticide target location (Kliot and Ghanim 2012; Yahouédo et al. 2017) they did not consider the possibility of soft sweeps, and thus may have missed additional targets of selection. Here we identified both shared and population-specific sweep windows that were almost entirely classified as soft sweeps (96.8% and 93.6%, respectively). Within these windows, we identified 22 genes with either well-characterized or putative roles in IR. To the best of our knowledge, several of our highlighted IR candidates (e.g., *ankyrin repeat domain-containing protein 29*, *split ends*, *sodium/potassium/calcium exchanger Nckx30C, muscle calcium channel subunit alpha-1*, or *potassium voltage-gated channel subfamily KQT member 1*), have not been reported as IR genes in *Ae. aegypti* and thus represent novel targets for functional validation. Interestingly, of our 13 shared selective sweep windows, only two of them had corresponding CLR peaks (defined by a CLR value > 200) and in our population-specific dataset, only 22% had corresponding CLR peaks. These results make sense given that CLR is based on a hard selective sweep model and the majority of our selective sweeps were classified as soft, further highlighting the need for methods specifically designed to detect the signatures of soft selective sweeps in order to more fully appreciate the landscape of adaptive evolution.

Although we were able to identify several novel IR candidate regions by searching for selective sweeps, we note that IR is a polygenic trait. One might therefore expect polygenic selection, in which the selected phenotype moves towards its optimum through the combined effect of small shifts in the frequencies of a large number of small-effect alleles (Berg and Coop 2014; Stephan 2016; Höllinger et al. 2019; Hayward and Sella 2022), to be the primary mode of adaptation in *Ae. aegypti*. Indeed, polygenic adaptation has been implicated in the evolution of complex traits, including insecticide resistance (Chen et al. 2023; Hobbs et al. 2023). However, selective sweeps may still occur during a polygenic shift to a new fitness optimum (Thornton 2019). Indeed, this may explain our success in finding sweep candidates at both known and potentially novel IR loci, even though diploS/HIC would be underpowered to detect more subtle allele frequency shifts that may also be contributing to adaptation for IR and other traits in *Ae. aegypti*.

While our examination of candidate sweep regions primarily focuses on genes associated with IR, we note that other traits may also be subject to recent positive selection. Given *Ae. aegypti’s* broad and expanding geographic distribution, robust invasive capacity, and the impact of ongoing climate change, it is highly likely some of the sweeps reported here are unrelated to IR. This is particularly relevant for the population samples from Brazil and Senegal, which are human specialists unlike the Kenya and Gabon samples which exhibit the ancestral generalist feeding behavior. The Senegal and Brazil samples thus have experienced additional selective pressures associated with domestication (e.g., adapting to urban environments and acquiring a preference for feeding on human blood; (Rose et al. 2020; Lozada-Chávez et al. 2025)). These reasons also likely underscore why our enrichment analysis did not yield significant results for previously known IR genes. It is also important to note that the list of 157 IR genes that we used to test for enrichment within sweep windows only represents a subset of the genes likely involved in IR and is therefore not comprehensive. Thus, we cannot rule out the possibility that an examination of a more complete set of IR genes in *Ae. aegypti* would reveal significant enrichment within sweep windows. Nonetheless, we concentrate our discussion on sweep windows containing selection on IR genes for two main reasons: 1) selection on these genes may impact mosquito control efforts, and 2) focusing on candidate regions with IR genes may help facilitate the difficult task of identifying the targets of selection: the fact that exceptionally strong signatures of selection have previously been observed near IR genes, suggests that, in sweep regions spanning multiple genes, it is reasonable to prioritize any IR genes as the most likely target of selection. Finally, although we do not analyze non-IR genes in this work, we provide a comprehensive list of all sweep calls with ≥ 0.95 confidence in any population sample in Supplemental Tables 3 and 4.

In this study we leveraged whole-genome data from multiple *Ae. aegypti* population samples together with a powerful machine learning tool to enrich our understanding of the genomic targets and mode of positive selection in this important vector species. In summary, our findings suggest that the rapid adaptation of *Ae. aegypti* to insecticides and other control strategies may, in part, be driven by soft selective sweeps. The persistence of high genetic diversity in out-of-Africa populations suggests that these population samples are likely to retain substantial adaptive potential, enabling rapid responses to future selective pressures. This underscores the importance of considering modes of adaptation beyond classic hard sweeps, as approaches that primarily target hard sweep signatures may overlook many adaptive variants. Our work highlights the need for systematic methods to detect soft sweeps and a more complete understanding of how standing genetic variation and polygenic selection contribute to adaptive evolution in insect vectors.

## METHODS

### Sampling, Sequencing, and Variant Calling

In this study, we used a previously curated whole genome sequencing dataset of 104 individuals from (Love et al. 2023). This dataset includes 18 samples from Santarém, Brazil; 13 from Franceville, Gabon; 19 from Kaya Bomu, Kenya; and 20 from Ngoye, Senegal, all taken from Rose et al. (2020), as well as 10 and 24 samples from Cali and Río Claro, Colombia, respectively, sequenced in Love et al. (2023). Detailed methods for DNA extractions, sequencing, and variant calling and filtering are available in (Love et al. 2023). Briefly, reads from all individuals were aligned to the AaegL5 reference genome (Matthews et al. 2018; NCBI accession GCF_002204515.2) using bwa-mem2 v. 2.1 (Vasimuddin et al. 2019), indels were removed and SNPs were filtered following GATK’s best practices (Koboldt 2020). Repetitive regions, non-biallelic SNPs, and non-uniquely mappable regions were removed and genotypes with qualities less than 20 were masked.

### Using Classifiers to Detect Selective Sweeps

To detect selective sweeps, we used diploS/HIC, a supervised machine learning approach that has been shown to be robust to nonequilibrium demography and has the capacity to distinguish between hard and soft sweeps in unphased data (Schrider and Kern 2016; Kern and Schrider 2018). In short, diploS/HIC classifies individual genomic windows into five categories (hard sweep, soft sweep, hard-linked, soft-linked, or neutral) based on a vector of windowed and transformed summary statistics that are input into a convolutional neural network (CNN). While this classification approach allows for robust inference based on many features jointly, it requires training datasets that consist of examples known to belong to each class. Therefore, we generated the training data using discoal (Kern and Schrider 2016). diploS/HIC then subdivides each simulated region into a number of equally sized, adjacent subwindows; we used the default value of 11 subwindows. For hard and soft sweeps, we selected the location of selection from a uniform distribution within each of the 11 subwindows. Specifically, we conducted 3,000 simulations where selection occurred uniformly within the leftmost subwindow, another 3,000 for the second subwindow, and so on for all 11 subwindows.

We sought to train each population sample’s classifier on simulations under demographic models that provide a reasonable fit to that population sample’s genomic data. To this end, we used Kent et al. (2025)’s population size histories, which were estimated using SMC++ for each population sample examined here. To account for variation in mutation and recombination rates across the genome, we allowed these parameters to vary across our training replicates. We set our mutation rate (*μ*) to vary uniformly from 8.82 × 10^−10^ to 8.82 × 10^−9^, giving a mean value of 4.85 × 10^−9^ which matches the estimate from Rose et al. (2023). We drew the recombination rate (*r*) from an exponential distribution with mean 4.85 × 10^−9^ (with values greater than 3\**r* not allowed), choosing this mean value because the average recombination rate in *Ae. aegypti* appears to be similar to our mean value of *μ* (Matthews et al. 2018; Chen et al. 2022); note that because this is a truncated exponential distribution the true mean value of *r* will be somewhat lower than our mean parameter value.

For the simulations involving sweeps, there are several additional parameters whose distributions are unknown, like the strength and timing of selection. For these, we drew from a wide distribution to ensure that the range of parameters seen during training encompasses those likely encountered during downstream inference. Specifically, we drew the selection coefficient from a log-uniform distribution ranging from 0.005 to 0.05 and the time of fixation of the beneficial allele, from *U*(0, 0.001). For soft and soft-linked sweeps, we simulated selection on a previously neutral standing variant, drawing the frequency of the previously neutral allele at the onset of selection, (*f*_0_), from *U*(0, 0.05). In total, we generated 3,000 simulated replicates for each class for each population. Because our goal was to identify strong selective sweeps that affect a wide stretch of the chromosome, we sought to simulate 2.75 Mb regions, but the version of discoal available at the time these simulations were performed was unable to simulate such regions due to memory constraints. We therefore simulated 550 kb regions and decreased the selection coefficient to *U*(0.001, 0.01), resulting in simulated sweeps with the same ratio of *s*/*r* as we would have obtained from 2.75 Mb simulations with *s*∼*U*(0.005, 0.05). More recently, discoal has been updated to be more memory efficient. This allows simulation of 2.75 Mb genomic regions. These simulations are very time consuming, so rather than regenerating training and test data for each population sample, we decided to directly assess the extent to which our approach of using smaller windows with smaller selection coefficients impacts our classification accuracy. To this end, we generated an additional test set of simulations for the Kenyan population sample using 2.75 Mb regions under the same demographic and selective parameters (i.e., *U*[0, 0.05]) . We then applied the original diploS/HIC classifier that was trained on 550 kb simulations to these newly generated 2.75 Mb simulations and evaluated classifier performance using confusion matrices (Supplemental Figure 13). The resulting confusion matrices show that the classifier actually performs slightly better on the larger-window simulations compared to its performance on the original 550 kb simulated windows. This is probably a result of the increased number of polymorphisms in the larger windows, which decreases variance in our summary statistic calculations. In any case, these results indicate that the classifier generalizes well to the larger window size and that the change in window scale does not negatively affect classification accuracy.

As described above, in our real data we masked genomic positions that had genotype qualities below 20. To incorporate heterogeneity in data quality into our training/test data, for each simulated window, we randomly selected a corresponding window from our empirical dataset and masked the same sites in the simulated window that had been masked in the empirical window. This ensured that our masking procedure affected our simulated data in the same way as our real data. Feature vectors were then calculated for each simulation replicate. These features vectors measure the spatial patterns of a number of population genetic summary statistics, including π (Tajima 1983), *θ^*_ω_ (Watterson 1975), Tajima’s *D* (Tajima 1989), the number of distinct haplotypes, average haplotype homozygosity (also referred to as H_1_ by Garud et al. 2015), H_12_ and H_2_/H_1_ (Garud et al. 2015), Z_ns_ (Kelly 1997), and the maximum value of Kim and Nielsen’s ω (Kim and Nielsen 2004); note that the latter two statistics, which measure linkage disequilibrium, were based on calculations of Rogers and Huff’s estimator of LD for unphased data (Rogers and Huff 2009). In addition to these commonly used statistics, diploS/HIC also includes in its feature vector estimates of the variance, skewness, and kurtosis of the distribution of the number of pairwise differences for all pairs of individuals in the sample; these summaries are useful for detecting hard and soft sweeps and discriminating between them (Kern and Schrider 2018). diploS/HIC subdivides the simulated region into 11 adjacent windows and calculates each of these statistics in each window, dividing that value by the sum of values for that statistic across all 11 subwindows. If the smallest of these 11 values for a given statistic is less than 0, the value for this subwindow is increased to zero, and the value for each other subwindow is increased by the same amount (i.e., the absolute value of the smallest subwindow is added to each subwindow in such cases).

After computing summary statistics, for each population sample we used diploS/HIC’s makeTrainingSets command to combine our simulations into a training set of 13,500 examples: 2,700 each for the hard sweep class (consisting of simulations where a hard sweep occurred in the central subwindow), the hard-linked class (where a hard sweep occurred in any subwindow but the central one), the soft sweep class (where a soft sweep occurred in the central subwindow), the soft-linked class (where a soft sweep occurred in any other subwindow), and the neutral class (where no sweep occurred). Similarly, we constructed an independent test set of 1,500 examples, 300 of each of the five classes. We then used diploS/HIC to train a CNN classifier for each population, holding out 10% of the training simulations as a validation set used for early stopping during training—if 5 consecutive epochs of training yielded no improvement (defined as a decrease in the categorical cross entropy loss function of at least 0.001), then training terminated and the best-performing set of neural network weights up to that point was used for the final classifier. We then applied each population sample’s classifier to its corresponding test set and evaluated performance using confusion matrices, precision recall curves (PR), and receiver operating characteristic (ROC) curves.

After training our classifiers, we then calculated feature vectors on the genomic data from the corresponding *Ae. aegypti* population samples, again using 2.75 Mb windows (each subdivided into 11 adjacent 250 kb subwindows); this was done across the three chromosomes of the *Ae. aegypti* reference genome, while the smaller scaffolds were ignored. We next applied our classifiers to the genomic data and classified a total of 4,738 windows in each of our four population samples. Because diploS/HIC is designed to detect the changes in diversity summaries at varying recombination distances away from a sweep, it cannot produce accurate predictions in non-recombining regions of the genome. We therefore calculated weighted average recombination rates in each of our predicted windows (using data from (Matthews et al. 2018)) and removed any regions where the recombination rate was equal to zero. This resulted in 4,631 windows classified in each of our population samples. Finally, to exclude regions of low confidence, we removed regions where the fraction of sites that were masked according to the data filtering criteria described above exceeded 0.85, which resulted in 4,599 windows in each population sample except for Brazil which contained 4,598 windows. These filtered windows were used for all downstream analyses. We then used diploS/HIC’s posterior class membership probability estimates in order to experiment with four different thresholds: 0.80, 0.90, 0.95, and 0.99. For a given threshold, we required the sum of the windows’ hard and soft sweep posterior probabilities to be greater or equal to the threshold before labelling the window as a sweep, and all windows whose highest-probability class was hard or soft but for whom the sweep probability threshold was not met were treated as uncertain and thus were not assigned to a class. We plotted the intersections of selective sweep locations for all populations for a given threshold using the UpsetR package (Conway et al. 2017).

### Accounting for the Impact of Classification Errors on the Relative Numbers of Hard and Soft Sweeps

To investigate how false positive rates may impact our ratio of hard sweep to soft sweep calls, we calculated the expected number of false discoveries obtained at a given sweep probability threshold for a specified population sample by using the *q-*value obtained at that threshold. We then made the conservative assumption that all false sweep discoveries are classified as soft sweeps and reduced our expected number of soft sweeps accordingly. We then examined the confusion matrix for the corresponding probability threshold to obtain the fraction of hard sweeps that are misclassified as soft, further reducing our total number of expected soft sweeps—during this step we conservatively ignored the possibility of soft sweeps being misclassified as hard.

### Identifying Novel High-Confidence Sweep Candidates

To generate a list of high-confidence regions that are sweeping in one or more population samples, we imposed sweep probability cutoffs as described above (See section: ***Using Classifiers to Detect Selective Sweeps***). We also calculated false discovery *q*-values for each sweep candidate as follows: first, for each sweep in each population sample, we obtained the sweep probability score from diploS/HIC, treating it as our current threshold, and then counted the numbers of both real and simulated sweeps that were equal to or exceeded this threshold. We then obtained the false discovery rate (FDR) by comparing the estimated number of neutral simulations misclassified as a sweep to the total number of predicted sweeps. Once we obtained an FDR for each candidate sweep and its corresponding threshold, *q*-values were calculated as described in (Storey 2002). This procedure was done separately for each population sample.

Our set of high-confidence, shared sweep candidates was generated by ensuring that the combined posterior probabilities of a sweep was ≥ 0.95 in all populations that contained the sweep. Similarly, the set of high-confidence population-specific sweeps consisted of windows where a sweep was predicted with ≥ 0.95 probability in a single population sample. Visualization of sweep candidates was performed using the UCSC Genome Browser (Kent et al. 2002; Perez et al. 2025), along with custom tracks highlighting several population genetic summary statistics calculated by diploS/HIC and CLR scores (Nielsen 2005; DeGiorgio et al. 2016) generated in Love et al. (2023). When searching for potential targets of selection, we focused on genes that were either contained entirely within or partially overlapped the sweeping window, while noting that it is possible that in some cases selection may be acting on a regulatory region impacting the expression of a gene lying outside of this window.

### Enrichment Analysis

We first performed an enrichment analysis to test whether genes previously implicated in IR were overrepresented among genomic regions under selection. This was done using a list of 157 *Ae. aegypti* genes previously implicated in insecticide resistance, compiled by Love et al. (2023). For each population, we only considered sweep windows with a posterior probability ≥ 0.95. To test for enrichment, we performed a permutation-based analysis where we permuted the association between the sweep statistic values and their genomic positions 10,000 times. For each permutation, we recalculated the number of IR genes that overlapped with sweep windows. Gene windows were extended by 1 kb on either side to account for uncertainty around sweep locations and the locations of nearby regulatory elements. One-sided *p*-values were calculated as the proportion of permutations in which the number of overlapping IR genes was greater than or equal to that observed in the real data. Fold enrichment was calculated as the ratio between the observed and mean permutated number of IR genes overlapping sweep windows.

Next, we performed an additional enrichment test (https://github.com/SchriderLab/permEnrichmentTest) following that of Love et al. (2023) to look for overrepresented GO terms among genes overlapping selective sweep windows with a posterior probability ≥ 0.8. Importantly, this method tests for significantly enriched GO terms after controlling for linkage between genes. It does this by concatenating the three chromosomes into a circular sequence and rotating all sweep scores around the circle by a random interval. We ran this enrichment analysis using 10,000 permutations of the sweep window posterior probabilities and then asked which GO terms were enriched in the selective sweep set compared to the permuted set. Finally, genes were extended by 1 kb in either direction.

## ACKNOWLEDGEMENTS

We would like to thank members of the Schrider lab for useful discussion. This work was funded by NIH awards R01HG010774 and R35GM138286. DRM was funded by NIH award R35GM148244.

## DATA AVAILABILITY

All sequencing data used in this paper are publicly available (see Love et al. 2023). All code used for the analyses presented in this paper can be found at https://github.com/SchriderLab/Detecting_Selection_Aedes_Aegypti_2026.

