## Supplementary Information for "Soft selective sweeps predominate in the yellow fever mosquito *Aedes aegypti*"

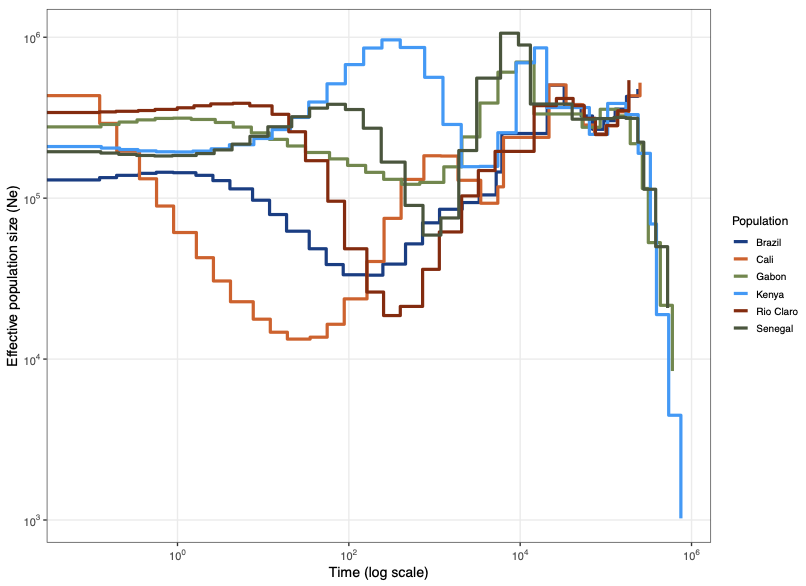

**Supplemental Figure 1.** Estimated population size (*N_e_*) history and split times inferred using SMC++ in Kent et al. 2025.

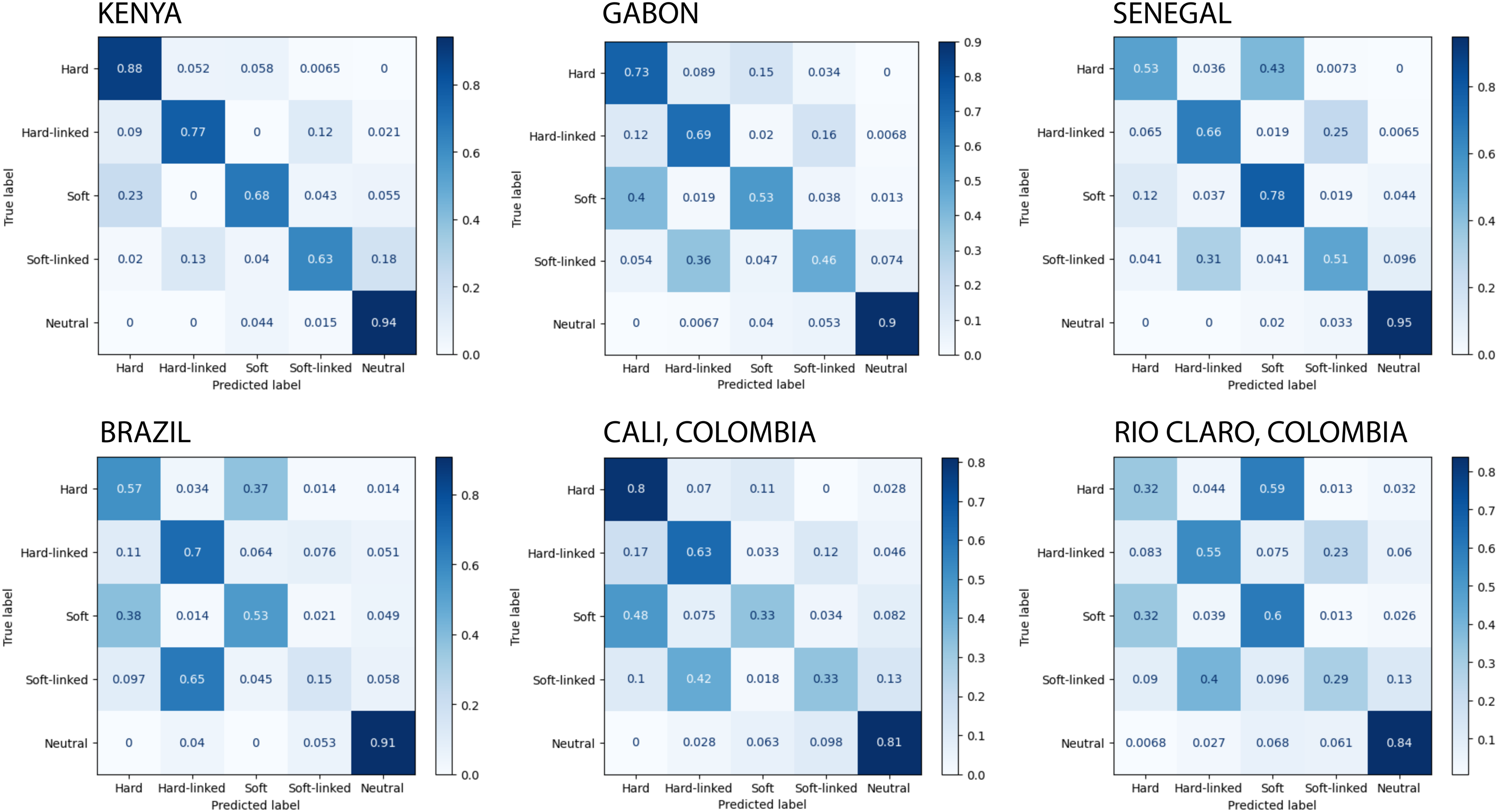
**Supplemental Figure 2.** Confusion matrices for each population sample with no posterior probability threshold applied.

**Supplemental Figure 3.** Confusion matrices for each population sample with a 0.80 posterior probability threshold applied.

**Supplemental Figure 4.** Confusion matrices for each population sample with a 0.90 posterior probability threshold applied.

**Supplemental Figure 5.** Confusion matrices for each population sample with a 0.95 posterior probability threshold applied.

**Supplemental Figure 6.** Confusion matrices for each population sample with a 0.99 posterior probability threshold applied.

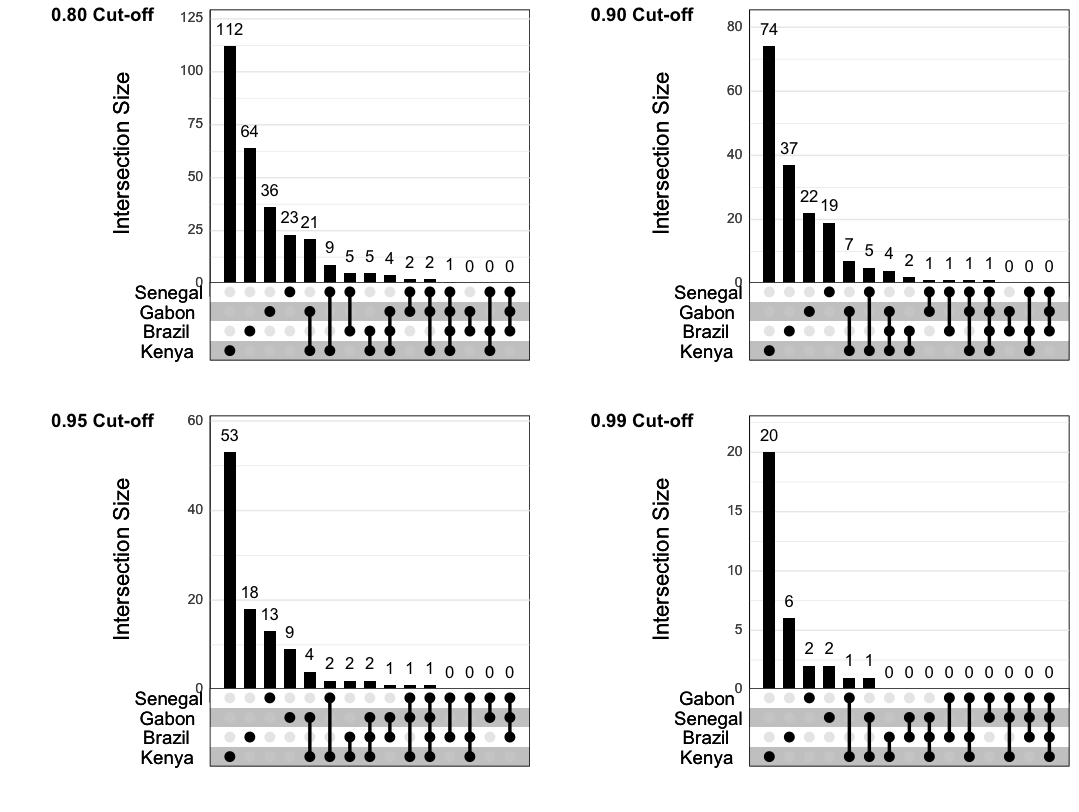

**Supplemental Figure 7.** Upset plots showing the overlap of sweeping windows discovered in diploS/HIC in each population sample at four posterior probability threshold cutoffs.

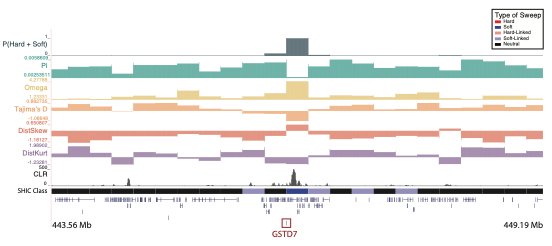
**Supplemental Figure 8.** A soft sweep in Brazil at a glutathione S-transferase gene (*GSTD7*) on chromosome two. The diploS/HIC classification track shows the class with the highest posterior probability, with soft sweeps as dark blue, soft-linked regions as light blue, hard sweeps as red, hard-linked as light red, and neutrally evolving regions in black. Above the diploS/HIC classifications are a subset of the summary statistics used by the classifier.

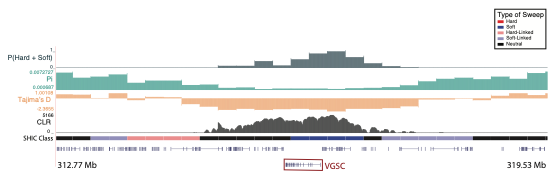

**Supplemental Figure 9.** A sweep in Brazil at a well-known IR gene: *VGSC* on chromosome three. The diploS/HIC classification track shows the class with the highest posterior probability, with soft sweeps as dark blue, soft-linked regions as light blue, hard sweeps as red, hard-linked as light red, and neutrally evolving regions in black. Above the diploS/HIC classifications are a subset of the summary statistics used by the classifier.

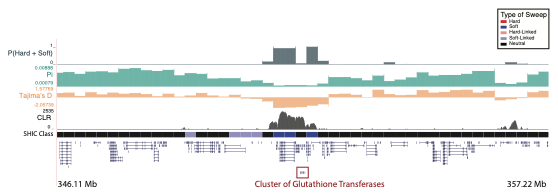
**Supplemental Figure 10.** A sweep in Brazil at a cluster of 15 glutathione S-transferase genes (*GST*) on chromosome two. The diploS/HIC classification track shows the class with the highest posterior probability, with soft sweeps as dark blue, soft-linked regions as light blue, hard sweeps as red, hard-linked as light red, and neutrally evolving regions in black. Above the diploS/HIC classifications are a subset of the summary statistics used by the classifier.

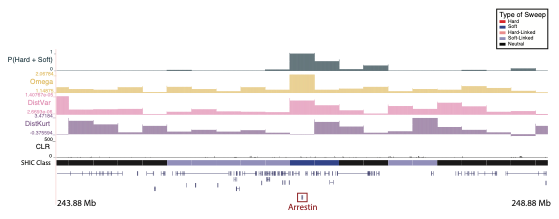
**Supplemental Figure 11.** A soft sweep in Kenya at an *arrestin* gene on chromosome two. The diploS/HIC classification track shows the class with the highest posterior probability, with soft sweeps as dark blue, soft-linked regions as light blue, hard sweeps as red, hard-linked as light red, and neutrally evolving regions in black. Above the diploS/HIC classifications are a subset of the summary statistics used by the classifier.

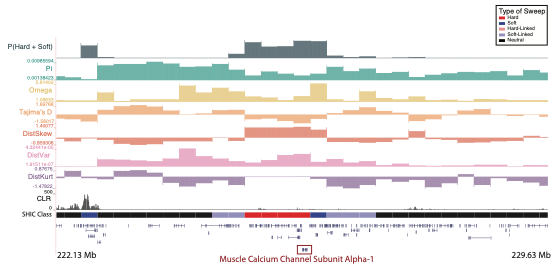

**Supplemental Figure 12.** A hard sweep in Brazil at a *muscle calcium channel subunit alpha-1* gene on chromosome two. The diploS/HIC classification track shows the class with the highest posterior probability, with soft sweeps as dark blue, soft-linked regions as light blue, hard sweeps as red, hard-linked as light red, and neutrally evolving regions in black. Above the diploS/HIC classifications are a subset of the summary statistics used by the classifier.

**Supplemental Figure 13.** Confusion matrices for the Kenyan population sample at four different posterior probability cutoffs. We generated an additional set of simulations using 2.75 Mb regions and applied the original diploS/HIC classifier (that was trained on the 550 kb simulations) to assess the extent to which our approach of using smaller windows with smaller selection coefficients impacts our classification accuracy. These results indicate that the classifier performs slightly better on the larger-window simulations.

**Supplemental Table 1:** To evaluate our finding that soft sweeps predominate, we applied a conservative approach to estimate the proportion of soft sweeps after accounting for 1) hard sweeps that may be misclassified as soft and 2) potentially false positive soft sweeps. It should be noted that this approach is conservative because we do not account for the possibility that hard sweeps are misclassified as soft sweeps.

**Supplemental Table 2:** For each window classified as a sweep, we calculated the false discovery rate for a given threshold (based on the summed probabilities of a window being a soft sweep and hard sweep) by comparing the number of misclassified neutral simulations to the total number of predicted sweeps. We then calculated each window's associated *q*-values based on the combined posterior probabilities and our predicted false positive rates obtained from simulations. We did this for all windows classified as a sweep and for each population sample individually.

**Supplemental Table 3:** The 13 sweep windows that are shared between 2-4 population samples with a posterior probability cutoff ≥ 0.95. The table includes chromosome, window start and end, the populations where the sweep is present, the type of sweep classified, and the genes containing or overlapping the window.

**Supplemental Table 4:** The 93 sweep windows that are population-specific sweeps with a posterior probability cutoff ≥0.95. The table includes chromosome, window start and end, the population where the sweep is present, the type of sweep classified, and the genes containing or overlapping the window.

**Supplemental Table 5:** Catalog of previously characterized (see Love et al., 2023) insecticide-resistance genes that was used to evaluate enrichment of IR genes within identified sweep windows.

**Supplemental Table 6:** Enrichment of known insecticide resistance genes within sweep windows across four *Aedes aegypti* population samples. For each population, the number of IR genes found within observed sweep windows (posterior probability cutoff ≥ 0.95) was compared with the mean number identified in 10,000 permuted sets of windows. No population sample shows statistically significant enrichment (all *P* > 0.05), although Senegal exhibits a higher but non-significant enrichment relative to expectation.

| **Population** | **Real IR Genes in Sweep Windows** | **Mean IR Genes in Permutated Windows** | **Fold Enrichment** | ***P*-value** |
| --- | --- | --- | --- | --- |
| **Brazil** | 1 | 0.8305 | 1.2 | 0.3529 |
| **Gabon** | 0 | 0.6257 | 0 | 1 |
| **Kenya** | 1 | 2.2594 | 0.44 | 0.6854 |
| **Senegal** | 3 | 0.6121 | 4.9 | 0.0701 |

**Supplemental Table 7:** Enrichment of GO terms within sweep windows across four *Aedes aegypti* population samples. Each sheet within this table contains the unique identifier for the GO term, the definition of the term, the number of significant genes in our dataset associated with this term, the average number of significant genes for this term across all permuted datasets (what would be expected by chance), the fold enrichment, the empirical *p*-value, and the false discovery rate (FDR) adjusted *p*-value.

**SUPPLEMENTARY TEXT**

*Population-Specific Sweep Windows Containing Well-Characterized Insecticide Resistance Genes*

We identified two high-confidence soft sweep windows in Kenya and one in Gabon that contained or overlapped three ATP-Binding Cassette (ABC) Transporter genes, ATP-binding cassette sub-family G member 4 (ABCG4; [chr2:455,500,001-455,750,000]), multidrug resistance-associated protein lethal (chr2:100,000,001-100,250,000) and multidrug resistance-associated protein 4 (chr3:138,250,001-138,500,000), respectively. In the first window, there is also a corresponding soft sweep in Brazil, although it did not meet the posterior probability cutoff at 0.86. Similarly, the second sweeping window also had a corresponding soft sweep in Gabon that did not meet the cutoff at 0.83. In humans, the multidrug resistant protein family (MRP) are transporters known for their role in causing multidrug resistance in tumor cells (Borst et al. 2000). In insects, ABC transporters have been documented as a group of detoxification-involved proteins (Labbé et al. 2011; Epis et al. 2014; Qi et al. 2016). In the mosquito *Culex pipiens pallens*, MDR1 and MDR2 were identified as proteins that were significantly enriched in cypermethrin resistance (Zhang et al. 2022) and in the Colorado potato beetle *Leptinotarsa decemlineata*, MRP4 was significantly upregulated in response to spinosa (Chen et al. 2023).

*Population-Specific Sweeps Windows with Putative or Emerging Insecticide Resistance Candidate Genes*

In Kenya, we found two sweeping windows that were classified as soft sweeps which overlapped or contained genes involved in the fatty acyl-CoA synthesis pathway, *acyl-CoA synthetase family member 4* (chr2:251,750,001-252,000,000) and *fatty acyl-CoA reductase 1* (chr3:309,250,001-309,500,000). *Acyl-CoA synthetase* (ACS), which activates fatty acids, was upregulated in the pea aphid (Cai et al. 2024) and was also found in a selection scan performed on *Anopheles gambiae*, along with several other IR genes (Dennis et al. 2024). It is also worth noting that this same window had a combined posterior probability of 0.94 in Senegal (marginally below our stringent 0.95 threshold). Fatty acyl-CoA reductases exhibit many biological functions, one of which being synthesis of cuticular hydrocarbons (CHCs; (Wang et al. 2024)). Many insecticides are absorbed through the insect cuticle and changes to the cuticle interface can slow the penetration of insecticides, thereby resulting in increased IR (Jacobs et al. 2023). In the cotton mealybug *Phenacoccus solenopsis*, a fatty acyl-CoA reductase (FAR) gene contributed to wax biosynthesis and RNA interference against *FAR* resulted in increased mortality post-deltamethrin treatments (Tong et al. 2022). Further, in a phosphine resistant strain of *Tribolium castaneum*, *FAR1* was upregulated and the cuticle structure was more continuous and tight than the susceptible strain (Kim et al. 2023).

We identified a window classified as a soft sweep in Gabon which contains the *membrane-associated progesterone receptor component 1* (chr1:15,500,001-15,750,000). This same window is classified as a soft sweep in Kenya and although the posterior probability of the window being a sweep (prob = 0.89) did not meet our cutoff, this may indicate a shared sweep between population samples. Membrane-associated progesterone receptor component 1 (PGRMC1) binds and stabilizes many different CYPs, thereby supporting P450 protein levels posttranscriptionally (McGuire et al. 2021). Further, when PGRMC1 was knocked out in mice livers, there was reduced enzyme levels of P450s. Although the role that PGRMC1 plays in insects is unclear, it was upregulated in response to imidacloprid exposure in honeybees (Kim et al. 2022).

In Kenya, we identified a sweeping window (chr1:48,750,001-49,000,000) that was classified as a soft sweep and overlapped the *D(2) dopamine receptor A*, which has been explored as new mode-of-action insecticide targets (Nuss et al. 2015). Further, pyrethroids are known to be potent releasers of dopamine (Elwan et al. 2006). In Brazil, we found a sweeping window that contains the *transcription factor grauzone* (chr1:63,750,001-64,000,000), is classified as a soft sweep, and contains a CLR peak > 540. The *transcription factor grauzone* is required for meiosis in oogenesis and it was significantly associated with resistance to pyrethroids in *Ae. aegypti* (Campbell et al. 2019) and *Anopeheles funestus* (Wondji et al. 2022).

In Brazil, we detected a window classified as a soft sweep which contained *NADH dehydrogenase 1 beta subcomplex subunit 5* (chr2:4,000,001-4,250,000). *NADH dehydrogenase 1 beta subcomplex subunit 5* encodes for a subunit of the mitochondrial NADH dehydrogenase (Complex 1) which has been shown to provide resistance to the toxicants paraquat and menadione in *D. melanogaster* (Gospodaryov et al. 2020). In another study, several subunits of the NADH dehydrogenase were upregulated in response to permethrin in *An. gambiae* which they attributed to a potential link between mitochondrial energy metabolism and detoxification (Vontas et al. 2005).
